# The right inferior frontal gyrus as pivotal node and effective regulator of the basal ganglia-thalamocortical response inhibition circuit

**DOI:** 10.1101/2022.05.26.493546

**Authors:** Qian Zhuang, Lei Qiao, Lei Xu, Shuxia Yao, Shuaiyu Chen, Xiaoxiao Zheng, Jialin Li, Meina Fu, Keshuang Li, Deniz Vatansever, Stefania Ferraro, Keith M. Kendrick, Benjamin Becker

**Author notes:** Corresponding authors: Benjamin Becker and Keith M. Kendrick No. 2006, Xiyuan Ave., West Hi-Tech Zone, Chengdu, Sichuan 611731, China. Qian Zhuang and Lei Qiao are joint first author.

## Abstract

The involvement of specific basal ganglia-thalamocortical circuits in response inhibition has been extensively mapped in the last few decades. However, the pivotal brain nodes and directed casual regulation within this inhibitory circuit in humans remains controversial. Here, we capitalize on recent progress in robust and biologically plausible directed causal modelling (DCM-PEB) and a large fMRI response inhibition dataset (n=218) to determine key nodes, their causal regulation and modulation via biological variables (sex) and inhibitory performance in the inhibitory control circuit encompassing the right inferior frontal gyrus (rIFG), caudate nucleus (rCau), globus pallidum (rGP) and thalamus (rThal).The entire neural circuit exhibited high intrinsic connectivity and an increasing rIFG inflow and its causal regulation over the rCau and rThal during response inhibition. In addition, sex and behavioral performance influenced the architecture of the regulatory circuits such that women displayed increased rThal self-inhibition and decreased rThal to GP modulation, while better inhibitory performance was associated with stronger rThal to rIFG communication. Furthermore, control analyses did not reveal a similar key communication in a left lateralized model. Together these findings indicate a pivotal role of the rIFG as input and causal regulator of subcortical response inhibition nodes.

## Introduction

Animal models and human neuroimaging studies convergently demonstrated that inhibitory control critically relies on highly specific basal ganglia-thalamocortical circuits (Alexander et al., 1986, 1991; Alexander and Crutcher, 1990; Aron et al., 2007; Jahfari et al., 2019; Morein-Zamir and Robbins, 2015; Pfeifer et al., 2022; Schall and Godlove, 2012; Stuphorn, 2015; Verbruggen and Logan, 2009; Wei and Wang, 2016). Dysregulations in this circuit have been implicated in disorders characterized by inhibitory control deficits, including addiction (Klugah-Brown et al., 2020; Morein-Zamir and Robbins, 2015; Zhou et al., 2018), attention deficit/hyperactivity (ADHD, Morein-Zamir et al., 2014; Sonuga-Barke, 2005), schizophrenia (Camchong et al., 2006; Feng et al., 2018; Mamah et al., 2007) and Parkinson Disorder (DeLong and Wichmann, 2015; Obeso et al., 2000). The key nodes within this response inhibition circuitry have been extensively mapped with convergent evidence suggesting critical contributions from the pre-supplementary motor area (pre-SMA) and lateral prefrontal cortex (lPFC), particularly inferior frontal gyrus (IFG, Aron et al., 2003; Dambacher et al., 2014; Hampshire et al., 2010; Maizey et al., 2020; Schaum et al., 2021; Verbruggen and Logan, 2008; Zhang et al., 2017) as well as striatal regions, in particular the caudate and putamen (Eagle et al., 2011; Ghahremani et al., 2012; Hampton et al., 2017; Kelly et al., 2004; Ott and Nieder, 2019; Robertson et al., 2015; Robbins, 2007).

Anatomical and neurochemical studies further suggest that response inhibitory control within this circuitry is modulated by dopaminergic and noradrenergic signaling (Bari et al., 2011; Ghahremani et al., 2012; Li et al., 2020; Pfeifer et al., 2022; Rae et al., 2016; Robertson et al., 2015). Dopamine receptor availability in the fronto-striatal circuits is significantly related to inhibition-related neural responses (Ghahremani et al., 2012; Pfeifer et al., 2022) and dopamine receptor availability in the lPFC modulates motor control via downstream regulatory projections to the striatum (Ott and Nieder, 2019; Vijayraghavan et al., 2016). Enhanced norepinephrine signaling facilitates response inhibition via modulation of the IFG and its connections with the striatum (Chamberlain et al., 2009; Rae et al., 2016), while the dorsal striatum represents an important locus of dopaminergic control of response inhibition (Ghahremani et al., 2012; Robertson et al., 2015) and the IFG plays an important role in top-down control of the basal ganglia regions (Buschman and Miller, 2014; Hampshire et al., 2010; Jahfari et al., 2012; Kim, 2014; Puiu et al., 2020; Renteria et al., 2018; Schaum et al., 2020; Tops and Boksem, 2011). In the basal ganglia-thalamocortical model of response inhibition (Alexander et al., 1986, 1991; Alexander and Crutcher, 1990) the thalamus relays information between the basal ganglia and cortex (Collins et al., 2018; Haber and Mcfarland, 2001; Haber and Calzavara, 2009; McFarland and Haber, 2002) - thus facilitating response inhibition and performance monitoring (Bosch-Bouju et al., 2013; Huang et al., 2018; Saalmann and Kastner, 2015; Tanaka and Kunimatsu, 2011) - via dense reciprocal connections with the basal ganglia and PFC (Guillery, 1995; Phillips et al., 2021; Xiao et al., 2009; Tanaka and Kunimatsu, 2011).

Convergent evidence from human lesion studies and neuroimaging meta-analyses demonstrates a right-lateralized inhibitory control network encompassing the right IFG (rIFG), right caudate nucleus (rCau), right globus pallidum (rGP) and right thalamus (rThal) (Aron et al., 2003; Chevrier et al., 2007; Garavan et al., 1999; Hung et al., 2018; Jahfari et al., 2011; Thompson et al., 2021). However, while extensive research has highlighted the critical role of these regions within a right-lateralized inhibitory control circuitry, the causal information flow and critical contribution of single nodes within this network have not been determined.

We therefore capitalized on a novel dynamic causal modelling (DCM) approach based on a priori specification of biologically and anatomically plausible models which allows estimation of directed causal influences between nodes and their modulation by changing task demands (Friston et al., 2003; Stephan et al., 2010) in the largest sample to-date (n=218). DCM further allows comparison of modulatory effective connectivity strength across different experimental conditions using Bayesian contrasts (Dijkstra et al., 2017) and in combination with the recently developed Parametrical Empirical Bayes (PEB) hierarchical framework (DCM-PEB method) allows modeling of both commonalities and differences in effective connectivity between subjects e.g. to determine the neurobiological basis of sex and behavioral performance variations (Friston et al., 2016; Zeidman et al., 2019a; Zeidman et al., 2019b).

To determine the causal information flow and critical nodes in the basal ganglia-thalamocortical circuits and whether these are modulated by biological factors (i.e. sex) and show functional relevance in terms of associations with performance we capitalized on DCM-PEB in combination with functional magnetic resonance imaging (fMRI) data collected in a large sample of healthy individuals (n=218) during a well-established response inhibition paradigm (emotional Go/NoGo task, see also Zhuang et al., 2021). To unravel the key nodes and causal influences within the inhibitory control network, we firstly estimated the effective connectivity between and within key regions involved in response inhibitory control within the rIFG-rCau-rGP-rThal functional circuit (right lateralized model) and secondly estimated sex differences and behavioral performance effects on connectivity parameters. To validate the hemispheric asymmetry of the inhibitory control network, an identical model of nodes was tested in the left hemisphere (left lateralized model).

Given convergent evidence on a pivotal role of the right IFG in mediating top-down cortical-subcortical control during response inhibition (Aron et al., 2003; Dambacher et al., 2014; Hampshire et al., 2010; Maizey et al., 2020), we predicted a greater modulatory effect on rIFG and its directed connectivity to both rCau and rThal in the NoGo compared to Go condition. Additionally, based on previous findings we expected a modulation of the key pathways by biological (i.e. sex, Li et al., 2006; Ribeiro et al., 2021; Sjoberg and Cole, 2018) and performance variations (Chang et al., 2020; Jahfari et al., 2011; Wei and Wang, 2016; Xu et al., 2016) with better response inhibition being associated with stronger causal regulation in the inhibition circuitry. Finally, we hypothesized a different causal structure for the left and right models given the hemispheric asymmetry in the inhibitory network (Aron et al., 2003; Chevrier et al., 2007; Hung et al., 2018; Jahfari et al., 2011; Thompson et al., 2021).

## Results

### Behavioral Results

The two-way repeated-measures ANOVA on accuracy found a significant main effect of inhibition (F(1,115)=21.73, p<0.001, η_p_^2^=0.16), with a higher accuracy for Go compared to No Go trials (Go trials: mean± SEM=98.47% ±0.31, No Go trials: mean ±SEM=70.34% ±1.44, Cohen’s d=2.48). No sex-differences were found for accuracy or reaction times (ps>0.18).

### BOLD Activation (GLM) Analysis

Examination of domain general inhibition (contrast: NoGo>Go) revealed a widespread fronto-parietal cortical and thalamo-striatal subcortical network including the IFG, striatal, pallidal and thalamic regions (**Figure 1 and Table 1**) during response inhibition. Group-level peaks in the rIFG, rCau, rGP and rThal were selected as centers of the ROIs for model testing (**Figure 2a**). No significant sex difference were observed in BOLD activation.

**Figure 1.**
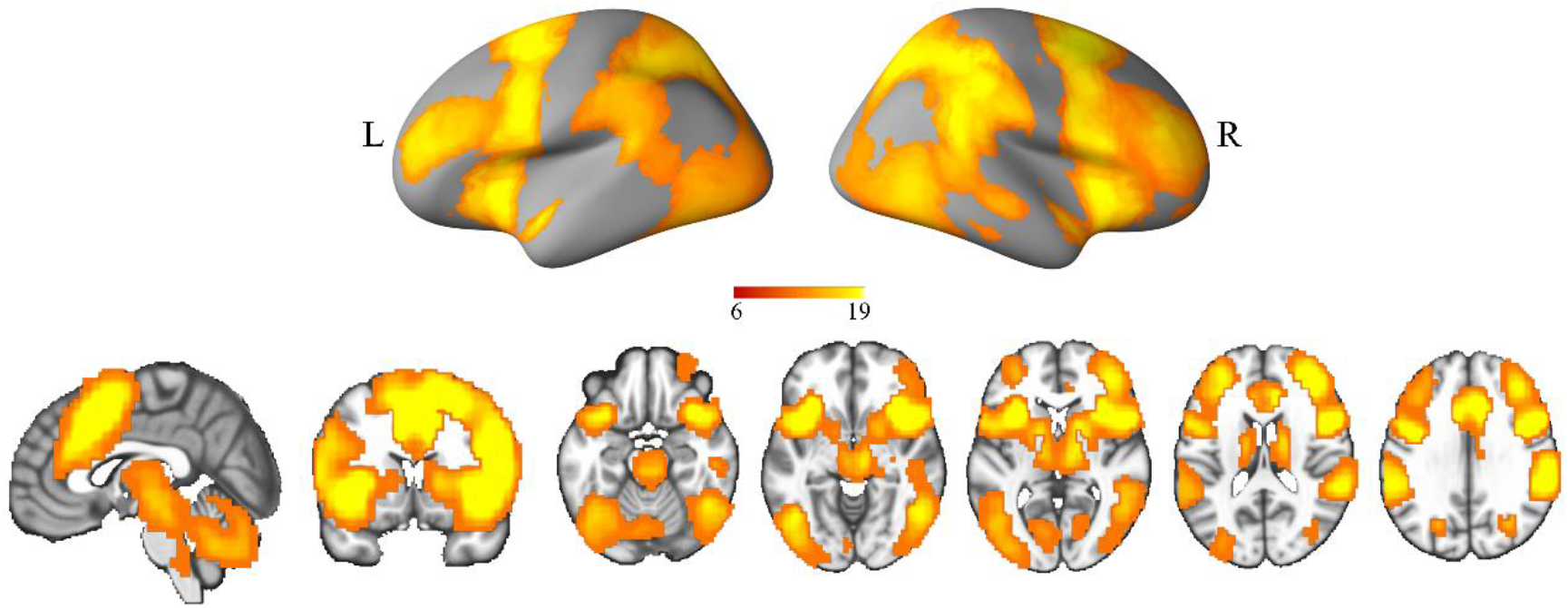
Brain activation maps for general response inhibition on whole brain level (contrast: NoGo > Go; p < 0.05 FWE, peak level). FWE, family-wise error; L, left; R, right.

**Figure 2.**
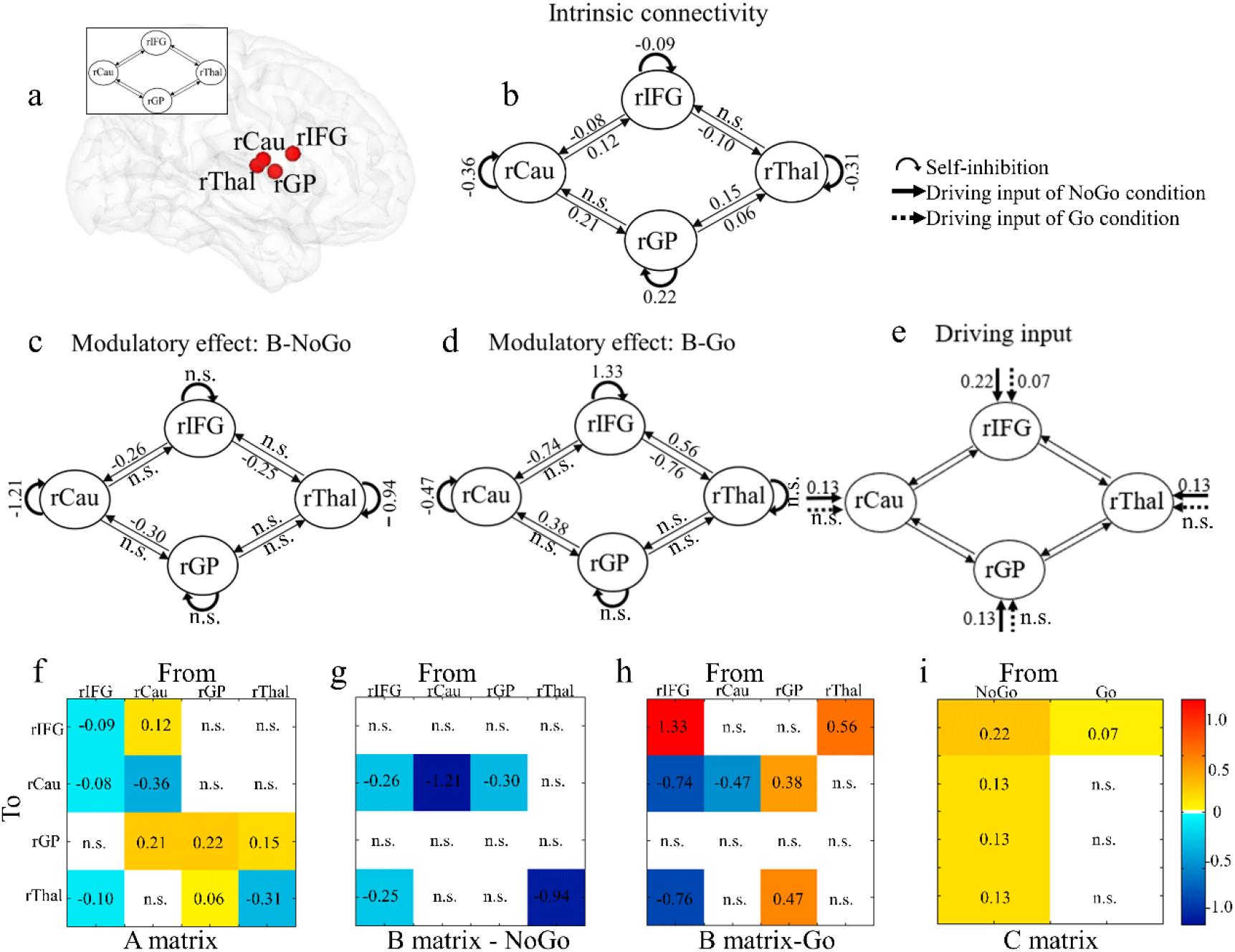
Location of regions included in the right model and group-level connectivity parameters. (a) Location of regions included in the right model. The A matrix: intrinsic connectivity across all experimental conditions (b, f). The B matrix: modulatory effect on effective connectivity between regions and self-inhibitions from NoGo (c, g) and Go condition (d, h). The C matrix: Driving inputs in ROIs in the NoGo and Go condition (e, i). Values in matrices reflect the connectivity parameters. Parameters with stronger evidence (posterior probability > 95%) are presented and subthreshold parameters are marked with “n.s.”.

**Table 1.** Activation and peak values for key regions included in the right model. Note: These clusters survived from the overlay between image masks of corresponding regions defined by Human Brainnetome Atlas ([84]) and group level brain activation maps (peak level, pFWE < 0.05) and thus served as regions of interest combined with the individual peak location search on the individual level. FWE, family-wise error; Cau, caudate nucleus; GP, global pallidum; IFG, inferior frontal gyrus; r, right; Thal, thalamus.

| Regions | Cluster K | Coordinates |  |  | t value |
| --- | --- | --- | --- | --- | --- |
|  |  | X | Y | Z |  |
| rIFG | 611 | 51 | 12 | 18 | 21.40 |
| rCau | 144 | 15 | -3 | 15 | 13.61 |
| rGP | 63 | 21 | 3 | 9 | 12.43 |
| rThal | 340 | 15 | -6 | 12 | 14.30 |

### Causal Connectivity (DCM) Analysis

For the A matrix, the diagonal cells represent self-connection which are unitless log scaling parameters and were multiplied with the default value of −0.5Hz (Zeidman et al., 2019a). Positive values indicate increased self-inhibition due to task condition and decreased responsivity to the inputs from the other regions of the network, while negative values indicate decreased self-inhibition and increased responsivity to the inputs from other nodes of the network (Zeidman et al., 2019a). Our findings revealed negative self-inhibition values for the rIFG, rCau and rThal but a positive value for the rGP (**Figure 2b, 2f**), indicating that the GP increased self-connection while the other nodes increased interaction with other nodes in the network.

For the off-diagonal cells in the A matrix, the values (in Hz) reflect the rate of change in the activity of the target region caused by the source region per second. Positive values reflect excitatory effects while negative values indicate inhibitory effects. In the forward direction (e.g. rIFG-rThal-rGP-rCau-rIFG), we found a significant negative connectivity from rIFG to rThal and positive connectivity from rThal to rGP as well as rCau to rIFG. In the backward direction (e.g. rIFG-rCau-rGP-rThal-rIFG), rIFG exhibited a negative inhibitory influence onto rCau, alongside an excitatory connection from rCau to rGP and rGP to rThal (**Figure 2b, 2f**). Although the connectivity from rThal to rIFG was not significant, a weak evidence (posterior probability=57%) for this connection was observed with a more lenient threshold.

Values in the B matrix represent the rate of change, in Hz, in the connectivity from source area to target area induced by the experimental conditions (Zeidman et al., 2019a). During inhibitory control (NoGo condition) the rIFG exerted a negative influence onto the rCau and rThal whereas the rGP exerted a negative influence on the rCau (**Figure 2c, 2g**). In addition, we found negative self-inhibition values in both rCau and rThal respectively. During the Go condition a negative influence of the rIFG on both rCau and rThal was observed (**Figure 2d, 2h**), while the positive influence was observed from the rGP to rCau and from rThal to rIFG. Moreover, we found a positive self-inhibition value in rIFG and a negative value in rCau. A Bayesian contrast (NoGo > Go) allowed us to compare the connectivity strength modulation during the different experimental conditions and revealed a very strong evidence (posterior probability >99%) that the causal influence of the rIFG to both, the rCau and rThal was stronger during inhibitory control (NoGo vs Go condition). This reflects that response inhibition critically requires a causal top-down cortical-subcortical regulation via the right IFG. We additionally found a very strong evidence (posterior probability >99%) for a considerably stronger inhibitory connectivity from rGP to rCau in the NoGo compared to Go condition.

The C matrix represents the rate of change in neural response of one brain region due to the driving input from an experimental condition (Zeidman et al., 2019a). During inhibitory control (NoGo) all regions (rIFG, rCau, rGP and rThal) exhibited excitatory driving input while during the Go condition only the rIFG exhibited excitatory input (**Figure 2e, 2i**). Bayesian contrasts directly comparing the conditions (NoGo > Go) demonstrated an increasing driving input specifically in the rIFG during engagement of cognitive control (NoGo > Go condition) with a 100% posterior probability.

### Sex Differences in Connectivity Parameters

Examining sex effects on intrinsic connectivity showed a negative influence from rThal to rGP in female compared to male subjects across all experimental conditions (**Figure 3a**). For the modulatory effects on connectivity, we found a greater self-inhibition in rThal in female than male subjects in the NoGo condition (**Figure 3b**). This suggests that for female subjects, rThal exhibits reduced sensitivity to inputs from the other regions of the selected network during response inhibition.

**Figure 3.**
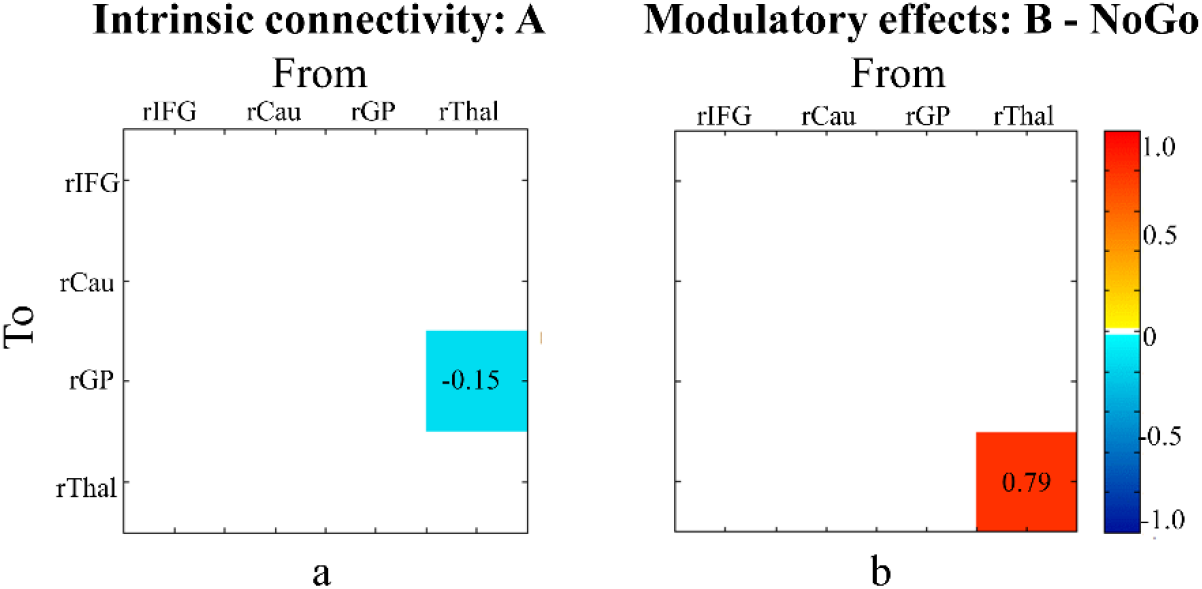
Sex effect on connectivity parameters in terms of A matrix and B matrix. (a) For intrinsic connectivity in A matrix, female subjects showed a more negative influence from rThal to rGP compared to male subjects. (b) In the NoGo condition, there is a greater self-inhibition in rThal in female than male subjects in terms of B matrix. Parameters with stronger evidence (posterior probability > 95%) are presented.

### Brain Behavior Associations: Inhibitory Behavioral Performance and Connectivity Parameters

Examining associations between inhibitory performance on the behavioral level (NoGo performance) and connectivity parameters revealed very strong evidence (posterior probability > 99%) that NoGo accuracy was positively associated with the directed connectivity from rThal to rIFG.

### DCM Analyses in the Left Hemisphere

To further validate the hemispheric asymmetry of the inhibitory control network, an identical model for the left hemisphere including lIFG, lCau, lGP and lThal was tested. In contrast to the right model, no directed influences from IFG to subcortical regions were observed in terms of matrix A in the left model, and the hemispheric models different in terms of inhibition induced connectivity changes and differed in terms of the driving inputs. The different causal structure in the left and right model indicated a hemispheric asymmetry in the inhibition network (details see **Supplementary Materials Figure S3**). Additional Bayesian analyses confirmed the lack of a robust cortical-subcortical pathway in the left hemisphere (**Supplementary Material**).

## Discussion

We capitalized on a combination of recent progress in biologically plausible causal hierarchical modelling (DCM-PEB) and a comparably large fMRI response inhibition dataset to determine causal information flow and key nodes within the extensively described basal ganglia-thalamocortical response inhibition circuits (Alexander et al., 1986, 1991; Alexander and Crutcher, 1990; Aron et al., 2007; Jahfari et al., 2019; Morein-Zamir and Robbins, 2015; Pfeifer et al., 2022; Schall and Godlove, 2012; Stuphorn, 2015; Verbruggen and Logan, 2009; Wei and Wang, 2016). Our neurocomputational model successfully validated a right-lateralized inhibitory control causal circuit and the best model showed significant intrinsic connectivity within this functional loop and captured an increasing causal influence of the cortical rIFG node on both the rCau and rThal as well as from the rGP to the rCau during inhibition. Direct comparison between different experimental conditions (e.g. NoGo and Go) revealed enhanced input into rIFG in terms of matrix C and increased connectivity from rIFG to rCau and rThal in the NoGo compared to the Go condition in terms of matrix B, suggesting a higher engagement of causal top-down cortical-to-subcortical control via the rIFG during inhibitory control. Although no sex differences were observed in inhibitory performance or BOLD activation, females exhibited decreased intrinsic connectivity from rThal to rGP and increased self-inhibition in rThal during the NoGo condition as compared to males. This indicates that similar behavioral performance in response inhibition might be mediated by different brain processes in men and women, particularly in thalamic loops. Moreover, a higher NoGo response accuracy was associated with stronger causal information flow from the rThal to rIFG in the NoGo condition, suggesting a particular behavioral inhibitory relevance of this pathway. Finally, our findings showed different left and right model structures, suggesting a hemispheric asymmetry in the inhibitory control network and confirming a critical role of the rIFG in implementing response inhibition. Together these findings identified a pivotal role of the rIFG and its effective connectivity with the rCau/rThal within the basal ganglia-thalamocortical circuit during response inhibition.

Causal modelling successfully determined a right lateralized inhibitory control causal circuit encompassing the rIFG, rCau, rGP and rThal (Aron et al., 2003; Chevrier et al., 2007; Hung et al., 2018; Jahfari et al., 2011; Thompson et al., 2021). In terms of the A matrix, a significant rIFG-rCau-rGP-rThal loop was observed with rIFG exhibiting a negative influence onto rThal, alongside a positive information flow from from rThal to rGP and rCau to rIFG in the forward direction. In the backward direction, we found significant negative connectivity from rIFG to rCau and positive connectivity from rCau to rGP as well as rGP to rThal. A more lenient threshold additionally revealed rThal to rIFG connections (posterior probability = 57%). Importantly, accounting for behavioral task context revealed a significant positive modulatory effect on rIFG in both NoGo and Go condition in terms of matrix C which was considerably stronger during response inhibition. The direct driving inputs into the rIFG are in line with its role in top-down target detection and attentional control in the context of response inhibition (Hampshire et al., 2010; Krämer et al., 2013) and indicate that the rIFG represents the key regulator of other nodes. In line with this hypothesis the best model in terms of matrix B revealed strong evidence for causal effective connectivity from the rIFG to both rCau and rThal during response inhibition (posterior probability >95%). This inhibitory pathway is consistent with previous reports on negative coupling between the rIFG and striatal regions during behavior control (Behan et al., 2015; Diekhof and Gruber, 2010). Notably, direct comparison using Bayesian contrast revealed a very strong evidence (posterior probability >99%) for increased modulatory connectivity from rIFG to rCau and rThal in the NoGo condition compared to the Go condition, suggesting the rIFG driven engagement of cortical-to-subcortical top-down control during response inhibition. Previous animal models and human neuroimaging meta-analyses have consistently identified the rIFG, as a key region implicated in dopaminergic and noradrenergic modulated inhibitory regulation (Bari et al., 2011; Hauber, 2010; Ott and Nieder, 2019; Pfeifer et al., 2022; Terra et al., 2020; Vijayraghavan et al., 2016; Zhukovsky et al., 2021) in particular during motor control and inhibition (Aron et al., 2003; Chamberlain and Sahakian, 2007; Puiu et al., 2020; Xu et al., 2016), while both, fronto-striatal and fronto-thalamic projections have been extensively involved in response inhibition (Ahissar and Oram, 2015; Bosch-Bouju et al., 2013; Marzinzik et al., 2008; Phillips et al., 2021; Schmitt et al., 2017; Sommer, 2003; Tanaka and Kunimatsu, 2011).

In addition to the cortical-subcortical pathways significant excitatory connectivity was observed from the rGP to rCau during the Go condition and switched to inhibitory connectivity when response inhibition was required during the NoGo condition. Direct comparison confirmed a considerably stronger inhibitory influence of the rGP on the rCau during response inhibition (posterior probability >99%), suggesting that communication between basal ganglia nodes is crucial for context-appropriate behavioral response control. The involvement of this pathway is in line with extensive neurophysiological evidence showing that GABA inhibitory projections from the external segment of the GP to the striatum play an essential role in cancelling a planned response when it is inappropriate (Mallet et al., 2016; Wei and Wang, 2016) (but see also subthalamic nucleus to substantia nigra pars reticulata pathways in Hikosaka et al., 2006; Mallet et al., 2016).

With respect to sex differences we observed that females exhibited decreased intrinsic connectivity from rThal to rGP and increased modulation by NoGo condition on self-inhibition in rThal compared to male subjects in the absence of performance differences. While previous findings on sex differences in response inhibition remained inconsistent (Chung et al., 2020; Gaillard et al., 2020; Gaillard et al., 2021; Li et al., 2006; Ribeiro et al., 2021; Sjoberg and Cole, 2018) the present findings suggest that our model was sensitive to biological variables and that separable information processes may underly response inhibition in men and women (see also Chung et al., 2020; Li et al., 2006). The functional relevance of the identified pathways was further underscored by a significant association between response inhibition performance and the causal influence from the rThal to rIFG in the NoGo condition demonstrating that this pathway involved in motor inhibition critically mediates behavioral success during inhibition (Wei and Wang, 2016).

Finally, our modelling tests confirmed a hemispheric asymmetry and support the critical role of right IFG circuit in response inhibition (Hung et al., 2018; Jahfari et al., 2011; Maizey et al., 2020). The different causal structures suggest a strong cortical-subcortical intrinsic connectivity and rIFG control on the right side. The left model revealed a different causal structure and null hypothesis tests showed moderate evidence for the difference between NoGo and Go condition’s modulatory effects on effective connectivity from lIFG to lCau and to rThal (e.g. lIFG to lCau: Bayes factor = 5.47; lIFG to lThal: Bayes factor = 8.20).

Response inhibition impairments have been observed in several disorders and identification of the rIFG as critical input and top-down regulator for response inhibition opens new targets for regional or connectivity-based neuromodulation such as real-time neurofeedback which has been established for these regions (Li et al., 2019; Weiss et al., 2022; Zhao et al., 2019). For instance, rIFG and response inhibition deficits have been determined in ADHD (Clark et al., 2007; Morein-Zamir et al., 2014) and targeting the rIFG in ADHD may be a promising treatment.

There are several limitations in the current study. First, in line with our main aim we did not account for emotional valence in the DCM model which may affect response inhibition(Schimmack and Derryberry, 2005). Second, we focused on specific nodes that were based on established basal ganglia-thalamocortical circuits proposed by Alexander (Alexander et al., 1986, 1991; Alexander and Crutcher, 1990) (see also neuroimaging meta-analysis (Hung et al., 2018). Other regions such as the STN (Aron et al., 2016; Aron and Poldrack, 2006; Chen et al., 2020) could be integrated in future studies.

In conclusion, our findings demonstrated a critical role of the rIFG as well as top-down cortical-subcortical control from the rIFG to rCau and rThal in response inhibition. The nodes and pathways of the model were sensitive to biological and performance variations. The nodes and pathways may represent promising targets to improve response inhibition in mental disorders.

## Materials and Methods

### Participants

N=250 healthy right-handed participants were enrolled in the current study and underwent a validated Go/NoGo fMRI paradigm. The data has been previously used to examine undirected functional connectivity within domain-general and emotion-specific inhibitory brain systems (Zhuang et al., 2021) and was part of larger neuroimaging project examining pain empathy (Li et al., 2018; Zhou et al., 2020), emotional face memory (Liu et al., 2022) and mirror neuron processing (Xu et al., 2022). After quality assessment n=218 subjects were included (104 males, **Supplementary Materials**). The study was approved by the local ethics committee and in accordance with the latest version of the Declaration of Helsinki.

### Response Inhibition Paradigm

A validated mixed event-related block design linguistic emotional Go/NoGo fMRI paradigm was employed (Goldstein et al., 2007; Protopopescu et al., 2005, details see Zhuang et al., 2021). Participants were required to make responses as accurately and quickly as possible based on orthographical cues, i.e. words were presented in normal or italic font. For normal font words subjects were instructed to perform a button-press (Go trials), while inhibiting their response to words presented in italic font (NoGo trials). Positive, negative and neutral words were included, however, given that the present study aimed to examine the causal influence within the general inhibition network and to increase statistical power in this respect the different emotional contexts were not further accounted for in the DCM analysis. Stimuli were presented in 2 runs and each run included 12 blocks (6 blocks: Go; 6 blocks: NoGo). Each Go block encompassed 18 normal font words (100% Go trials) while each NoGo block encompassed 12 normal font words (66.7% Go trials) and 6 italicized font words (33.3% NoGo trials). Further details in (Zhuang et al., 2021) and **Supplementary Materials**.

### Behavioral Data Analysis

In our previous study we demonstrated that subjects exhibited more errors during inhibitory control (i.e., NoGo>Go) as well as faster responses in positive Go contexts and lower accuracy in positive NoGo contexts (Zhuang et al., 2021). Given that sex-differences were examined in the DCM model the present analyses additionally examined sex-differences on accuracy and reaction times (**Supplementary Materials)**. Given age-related effects on inhibition (Rey-Mermet et al., 2018; Rubia et al., 2007) age was included as covariate.

### MRI Data Acquisition and Preprocessing

MRI data were collected on a 3T MRI system using standard sequences and were initially preprocessed using validated protocols in SPM 12 (details see **Supplementary Materials**)

### GLM Analysis

An event-related general linear model (GLM) was established in SPM12. To examine domain general inhibitory control (irrespective of emotional context) the overarching inhibitory control contrast was modelled (e.g. all NoGo>all Go trials) and convolved with the canonical hemodynamic response function (HRF). Six head motion parameters were included in the design matrix to control movement-related artifacts and a high-pass filter (1/128Hz) was applied to remove low frequency components. The contrast of interest (contrast: NoGo>Go) was created and subjected to one-sample t-test at the second level. In line with previous studies (Aron et al., 2003; Chevrier et al., 2007; Hung et al., 2018; Jahfari et al., 2011; Thompson et al., 2021), group-level (contrast: NoGo>Go) peaks in the IFG, Cau, GP and Thal within the identified general inhibition network were then used to define individual-specific regions of interest (ROIs) for the DCM analysis. Additionally, a two-sample t-test was conducted (contrast: NoGo>Go) to examine sex-dependent effects on the response inhibition network. Analyses were corrected for multiple comparisons using a conservative peak-level threshold on the whole brain level (p<0.05 family-wise error, FWE).

### Dynamic Causal Modeling and Node Definition

A DCM analysis was employed to determine directed causal influences according to the circuitry model proposed by Alexander et al. (Alexander et al., 1986, 1991; Alexander and Crutcher, 1990). The DCM approach allows to construct a realistic neuronal model of interacting regions and to predict the underlying neuronal activity from the measured hemodynamic response (Friston et al., 2003; Stephan et al., 2007). To this end directed causal influences between the key regions including IFG, Cau, GP and Thal in the basal ganglia-thalamocortical loop and their modulation via experimental manipulations (engagement of motor inhibitory control) were examined. In line with previous neuroimaging studies and meta-analyses demonstrating a right-lateralized inhibition model (right model) encompassing the rIFG, rCau, rGP, rThal (Aron et al., 2003; Chevrier et al., 2007; Hung et al., 2018; Jahfari et al., 2011; Thompson et al., 2021) our main hypothesis testing focused on the right lateralized network. To further validate the hemispheric asymmetry of the inhibitory control network an identical model was tested for the left hemisphere including the lIFG, lCau, lGP, and lThal. In line with previous studies, we combined atlas-based masks (Human Brainnetome Atlas, Fan et al., 2016) with group-level and individual level activity maps to generate the corresponding nodes (Fernández-Espejo et al., 2015; Holmes et al., 2021; Qiao et al., 2020; Van Overwalle et al., 2020).

### Model Specification and Estimation

A two-step DCM analysis was performed using the DCM-parametric empirical Bayes (PEB) approach (Zeidman et al., 2019a; Zeidman et al., 2019b). On the first-level, time-series from four ROIs (rIFG, rCau, rGP, rThal) were extracted. A full DCM model was specified for each subject and all connectivity parameters in both forward (e.g. rIFG-rThal-rGP-rCau-rIFG) and backward (e.g. rIFG-rCau-rGP-rThal-rIFG) directions were estimated. We estimated three key DCM parameters: (1) the A matrix reflecting all connections including forward and backward connectivity between ROIs and self-inhibitions in each ROI, (2) the B matrix representing modulatory effects of Go and NoGo condition on all connections, (3) the C matrix representing the driving inputs into ROIs from Go and NoGo conditions separately. Given that all inputs in the model were mean-centered, intrinsic connectivity in the A matrix indicates mean effective connectivity independent of all experimental conditions. The model was estimated using Variational Laplace (Friston et al., 2007). Further details **Supplementary Material.** At the second (group) level, we constructed a PEB model over the first-level estimated parameters. In accordance with previous studies (Bencivenga et al., 2021; Rupprechter et al., 2020), we evaluated the explained variance by the model on the individual level - higher values reflect better model inversion (Zeidman et al., 2019a) – and then we only included subjects with >10% of explained variance in the PEB model. A total of 118 subjects (56 males, age: mean ± SEM = 21.57 ± 0.21 years) were included for further analyses. The differences on behavioral performance were examined between the excluded and included subjects and no significant differences were found (all ps≥0.23, for details see **Supplementary Material**), suggesting no evidence of biased selection.

The primary aim of the present study was to establish a causal neurobiological model for response inhibition and to determine the interaction between key players in this circuitry. To evaluate the model three PEB analyses were carried out separately for A, B and C matrices. Separate analyses examined sex and performance variations (details see **Supplementary Material**).

Next, to identify the model that best represented our data, Bayesian Model Reduction (BMR) was performed to compare the free energy of the full model with numerous reduced models for which specific parameters were “switched off” (Friston et al., 2016). An automatic greedy search procedure (iterative procedure) was employed to facilitate an efficient comparison of thousands of models. In this procedure parameters which do not contribute to free energy were pruned away. Next, Bayesian Model Average (BMA), performing a weighted average of the parameters of each model, was calculated over the 256 models obtained from the final iteration (Friston et al., 2016).

Finally, to compare the effective connection strength, especially the cortical-subcortical connectivity and driving inputs into each region from different experimental conditions (NoGo and Go condition), Bayesian contrasts (Dijkstra et al., 2017) were computed over parameters from the B and C matrices. Group-level estimated parameters were thresholded at posterior probability > 95% (indicating strong evidence, Kass and Raftery, 1995) based on free energy.

## Acknowledgements

This work was supported by the by the National Key Research and Development Program of China (grant number: 2018YFA0701400 - BB), National Natural Science Foundation of China (grant numbers 31530032 – KMK, 91632117 - BB), Key Technological Projects of Guangdong Province (grant number 2018B030335001 – KMK).

## Author contributions

**Qian Zhuang:** Formal analysis; Investigation; Writing-original draft. **Lei Qiao**: Formal analysis; Validation. **Lei Xu**: Project administration. **Shuxia Yao**: Conceptualization; Supervision**. Shuaiyu Chen:** Formal analysis**. Xiaoxiao Zheng, Jialin Li, Meina Fu and Keshuang Li**: Project administration. **Deniz Vatansever and Stefania Ferraro:** Validation. **Keith M. Kendrick and Benjamin Becker:** Conceptualization; Funding acquisition; Project administration; Resources; Supervision; Validation; Writing.

## Declaration of Conflicting Interests

The authors declared no conflicts of interest with their research, authorship or the publication of this article.

## Notes

https://osf.io/h3mg2/

### Competing Interest Statement

The authors have declared no competing interest.

### Summary of Updates

The format of the manuscript has been revised according to publication's guideline

