## Supplement for "The right inferior frontal gyrus as pivotal node and effective regulator of the basal ganglia-thalamocortical response inhibition circuit" (from another version of this preprint: version 1, 2022-05-28)

Zhuang, Qiao et al.,

Contact:

### Participants and Quality Control

Data from 11 subjects were lost due to technical issues during fMRI (n=5) and behavioral data acquisition (n=6). Data from another 21 subjects were further excluded due to excessive head motion (>2.5mm or 2.5 degrees, n = 12) or an incomplete number of runs due to technical issues (incomplete data, n=9) leading to a sample of 218 subjects (104 males, age: mean ± SEM = 21.58 ± 0.15 years) for the final behavioral and fMRI data analyses.

### Stimuli and Paradigm

A total of 54 Chinese words (18 positive, 18 negative, 18 neutral), matched for length and frequency were included. While positive words described positive emotions, negative words described fearful and threatening emotions, and neutral words described neutral experiences. Given that the present study aimed to examine the causal influence within the general inhibition network, the different emotional contexts were not further accounted for in the DCM analysis. Stimuli were further evaluated by an independent sample of n=18 individuals who rated the emotional category (yes or no), intensity (1-9 scale) and imaginability (1-9 scale) of the word stimuli (details see **Table S1**). Stimuli were presented in 2 runs and each run included 12 blocks (6 blocks: Go; 6 blocks: NoGo). Each Go block encompassed 18 normal font words (100% Go trials) while each NoGo block encompassed 12 normal font words (66.7% Go trials) and 6 italicized font words (33.3% NoGo trials). Before each block, a 1400ms instruction was presented to indicate the Go/NoGo condition (see **Figure S1**). Each stimulus was presented 300ms interleaved by a 900ms inter-stimuli-interval (total block duration=21.6s), blocks were separate by a 16s inter-block-interval which served as low level baseline. The paradigm lasted 16min.

### MRI Data Acquisition

MRI data were collected using a 3T GE Discovery MR750 system (General Electric Medical System, Milwaukee, WI). 488 volumes of T2*-weighted echo planar images were acquired (acquisition parameters: repetition time, 2000ms; echo time, 30 ms; slices, 39; slice-thickness, 3.4mm; gap, 0.6mm; field of view, 240 × 240 $mm^2$; matrix size, 64 × 64; flip angle, 90°). To improve normalization of the functional images and identify individuals with apparent brain pathologies, high-resolution whole brain T1-weighted images were obtained using a 3D spoiled gradient echo pulse sequence (acquisition parameters: repetition time, 6ms; echo time, minimum; flip angle, 9°; field of view = 256 × 256mm; acquisition matrix, 256 × 256; thickness, 1mm; 156 slices). OptoActive MRI headphones (http://www.optoacoustics.com/) were used to reduce acoustic noise during MRI data acquisition.

### fMRI Data Preprocessing

Functional MRI data were preprocessed using SPM12 software (Wellcome Trust Center of

Neuroimaging, University College London, London, United Kingdom). The first 10 volumes for each run were deleted to allow magnet steady data. The remaining functional images were processed using standard preprocessing procedures including: slice-timing and head motion correction, spatial normalization to Montreal Neurological Institute (MNI) standard space (by means of co-registration to the T1-weighted structural images and the application of the transformation matrices obtained from the segmentation of the structural images), resampling with a 3 x 3 x 3 mm resolution, and spatial smoothing using a 8mm full-width at half-maximum (FWHM) Gaussian kernel.

**Behavioral Analyses**

Given that we intended to additionally explore sex-differences in the DCM analysis we also examined sex differences at the behavioral level. To this end a two-way repeated ANOVA with sex (male vs. female) as between-subject factor and inhibition (NoGo vs. Go) as within-subject factor was conducted for response accuracy. For reaction times on Go trials with correct responses, a two-sample t test with sex (male vs. female) as grouping variable was performed. Given previous findings showing age-related effects on inhibition control (Rey-Mermet et al., 2018; Rubia et al., 2007), age was included as covariate for both accuracy and reaction time analyses.

**Differences in Behavioral Performance between excluded and included subjects**

The behavioral performance differences between excluded and included groups in the right model analysis were examined by means of mixed ANOVAs. For accuracy, mixed ANOVA with inhibition (NoGo vs. Go) as within-subject factors, group (included vs. excluded) as between-subject factors were performed. For reaction time on corrected Go trials, a two-sample t test with group (included vs. excluded) as grouping variable was performed. Given sex and age's potential effect on behavioral inhibition control (Li et al., 2006; Rey-Mermet et al., 2018; Rubia et al., 2007), these two variables were included as covariates for both accuracy and reaction time analysis.

The mixed ANOVA on accuracy showed neither significant group-related main ($p = 0.74$) nor interaction effect ($p = 0.77$). For reaction time on corrected trials, the two-sample t test showed no group related difference ($p = 0.23$). These results indicate no differences between excluded and included groups on behavioral performance.

### DCM: Node Selection and Regions of Interest

In line with previous neuroimaging studies and meta-analyses demonstrating a right-lateralized inhibition model (right model) encompassing the rIFG, rCau, rGP, rThal (Aron et al., 2003; Chevrier et al., 2007; Hung et al., 2018; Jahfari et al., 2011; Thompson et al., 2021) our main hypothesis testing focused on the right lateralized network. To further validate the hemispheric asymmetry of the inhibitory control network an identical model was tested for the left hemisphere including the lIFG, lCau, lGP, and lThal. To account for individual variations in the exact location of the inhibitory control systems, and to increase the sensitivity of the analysis, subject-specific ROIs were created using 6mm spheres around the nearest local maxima within a spherical mask (16mm radius) centered at the group-level GLM activation peak (contrast: NoGo > Go) in the identified regions (IFG, Cau, GP, Thal) defined by Human Brainnetome Atlas (Fan et al., 2016). In line with previous studies (Fernández-Espejo et al., 2015; Qiao et al., 2020), the radius of the spherical mask was set to twice the smoothing kernel size (16mm). In case no voxels survived a threshold of $p < 0.05$ uncorrected, on the individual level a more lenient threshold ($p < 0.10$ or $p < 1.00$ uncorrected) was employed (for an identical approach see Holmes et al., 2021; Van Overwalle et al., 2020). Next, the first principal eigenvariate of time series for each ROI was extracted and adjusted for effects of interest from all experimental conditions (NoGo and Go conditions) using a GLM model in which six head movement parameters and other effects of no interest were regressed out.

**Model Specification and Estimation**

A two-step DCM analysis was performed using the DCM-parametric empirical Bayes (PEB) approach (Zeidman et al., 2019a; Zeidman et al., 2019b), which allows estimation of both averaged effective connectivity across subjects and effects of covariant on connectivity parameters at the group level using a hierarchical Bayesian model. This method assumes that all subjects have the same model architecture, and the connection strength differs due to task condition. On the first-level, time-series from four ROIs (rIFG, rCau, rGP, rThal) were extracted. A full DCM model was specified for each subject and all connectivity parameters in both forward (e.g. rIFG-rThal-rGP-rCau-rIFG ) and backward (e.g. rIFG-rCau-rGP-rThal-rIFG) directions were estimated. In line with previous studies (Friston et al., 2016; Zeidman et al., 2019a), we estimated three key DCM parameters: (1) the A matrix reflecting all connections including forward and backward connectivity between ROIs and self-inhibitions in each ROI, (2) the B matrix representing modulatory effects of Go and NoGo condition on all connections, (3) the C matrix representing the driving inputs into ROIs from Go and NoGo conditions separately. Given that all inputs in the model were mean-centered, intrinsic connectivity in the A matrix indicates mean effective connectivity independent of all experimental conditions.

The estimation of model used Variational Laplace (Friston et al., 2007), which aims to match the predicted time series with the observed time series with maximum accuracy and minimized differences between estimated parameters and their prior values. The variational free energy was obtained to qualify the model with a higher value indicating the best trade-off between model accuracy and complexity.

The primary aim of the present study was to establish a causal neurobiological model for response inhibition and to determine the interaction between key players in this circuitry. To evaluate the model three PEB analyses were carried out separately for A, B and C matrices. In each PEB design matrix, the first regressor represents a column of ones modelling the group mean connectivity. To further determine the modulation of the identified circuitry by biological and behavioral variations we additionally examined sex differences and performance modulations. To this end, we set sex and NoGo response accuracy (covariates of interests) as the second regressor in two different models to evaluate their impact on the connectivity parameters. While NoGo accuracy, Go response accuracy, Go reaction time and age were regressed out to evaluate sex differences, sex, Go response accuracy, Go reaction time and age were regressed out for NoGo performance effects on design matrices as nuisance variables (all of them were mean-centered). To further evaluate the robustness of sex and

NoGo performance modulation these factors were included as independent variables in the covariate matrix, and additional control analyses were performed including only a column of ones and covariates of interests (gender or NoGo performance) in the design matrix.

**Control Analyses**

To enhance confidence in the sex-dependent effects, additional control analyses with a column of ones and gender as covariates was conducted. Results showed stable sex differences in connectivity from rThal to rGP in intrinsic connectivity (**Figure S2.a**) and also increased self-inhibition in rThal in the NoGo condition for female subjects (**Figure S2.b**). For the performance effect, the result also remained stable over additional control analyses with a column of ones, NoGo performance and gender as covariates.

**Left Hemispheric DCM Model**

To further validate the hemispheric asymmetry of the inhibitory control network, an identical model for the left hemisphere including lIFG, lCau, lGP and lThal (**Figure S3.a**) was tested. Subjects with < 10% explained variance were excluded and 82 subjects (40 males, age: mean ± SEM = 21.24 ± 0.27 years) were included for the final DCM analyses. In contrast to the right model, no directed influences from IFG to subcortical regions were observed in terms of matrix A in the left model (**Figure S3.b, S3.f**). Although the results showed modulatory effects of NoGo and Go conditions on the connectivity from IFG to Cau and Thal in both left and right models, the modulation effect of experimental condition on GP to Cau connectivity was only found in the right model (**Figure S3.c, S3.d, S3.g, S3.h**). Additionally, the NoGo condition showed an inhibitory modulatory effect on the connectivity from Cau to GP in the left but not the right model and the Go condition showed an excitatory modulatory effect on the connectivity from Thal to IFG in the right but not the left one. Moreover, the two models had a similar pattern for the driving inputs of the NoGo condition on regions but not the Go condition (**Figure S3.e, S3.i**). The different causal structure in the left and right model indicated a hemispheric asymmetry in the inhibition network.

**Null Hypothesis Tests of Modulation Effects of Different Experimental Conditions on the Cortical-subcortical Effective Connectivity in the Left Model Analysis**

To further test the null hypotheses with respect to the differences between the modulatory effect of experimental conditions on the effective connectivity from lIFG to lCau and to lThal, Bayes analyses were conducted using JASP (version 0.14.1.0). Bayes analysis using the Bayesian paired-sample T tests found Bayes factors ($BF_{01}$) of 5.47 and 8.20 for directed connectivity from lIFG to lCau and to lThal, suggesting that non-significant hypotheses for the differences between modulatory effect of NoGo and Go conditions on cortical-subcortical effective connectivity were moderately fitted for the present data (Jeffreys, 1998). These results further confirm the lack of the cortical-subcortical pathway's engagement in the left response inhibition network.

**Table S1. Results from emotional valence, intensity, imagination rating as well as word frequency.**

| Measurements | Stimuli | | | F | p |
|---|---|---|---|---|---|
| | Positive | Negative | Neutral | | |
| Emotional valence rating | 94.58% ± 1.07% | 96.00% ± 1.07% | 96.21% ± 0.73% | 0.95 | 0.35 |
| Frequency | 1965.58 ± 340.04 | 1007.24 ± 446.22 | 1725.54 ± 288.05 | 1.87 | 0.16 |
| Intensity | 5.73 ± 0.17 | 6.15 ± 0.23 | | 1.47 (t) | 0.15 |
| Imagination | 5.76 ± 0.23 | 6.07 ± 0.22 | 5.70 ± 0.12 | 1.02 | 0.37 |

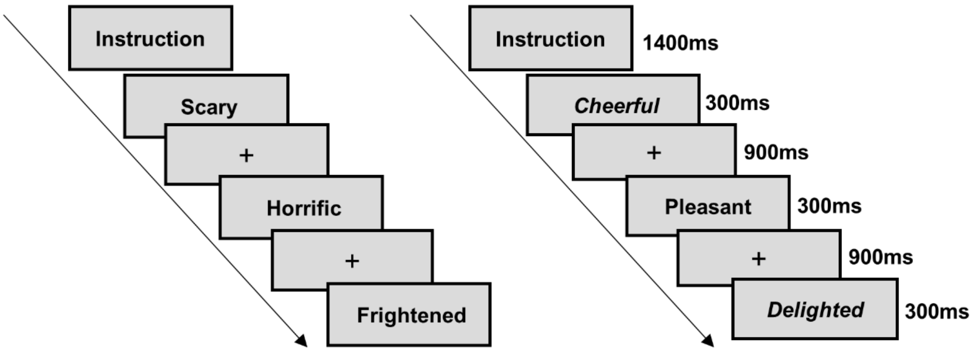


**Figure S1.** The emotional Go/NoGo paradigm. Timing for a negative Go block (left) and positive NoGo block (right) are depicted.

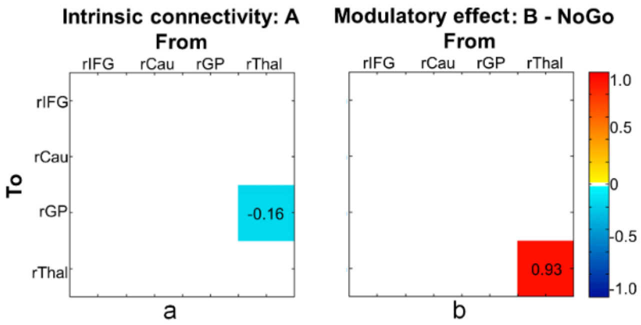


**Figure S2.** Additional control analyses for sex differences on connectivity parameters. (a) For intrinsic connectivity, female subjects showed a more negative influence from rThal to rGP compared to male subjects. (b) For modulatory effect on connections, there is an increased self-inhibition in rThal in female compared to male subjects in the NoGo condition. Parameters with stronger evidence (posterior probability > 95%) are presented.

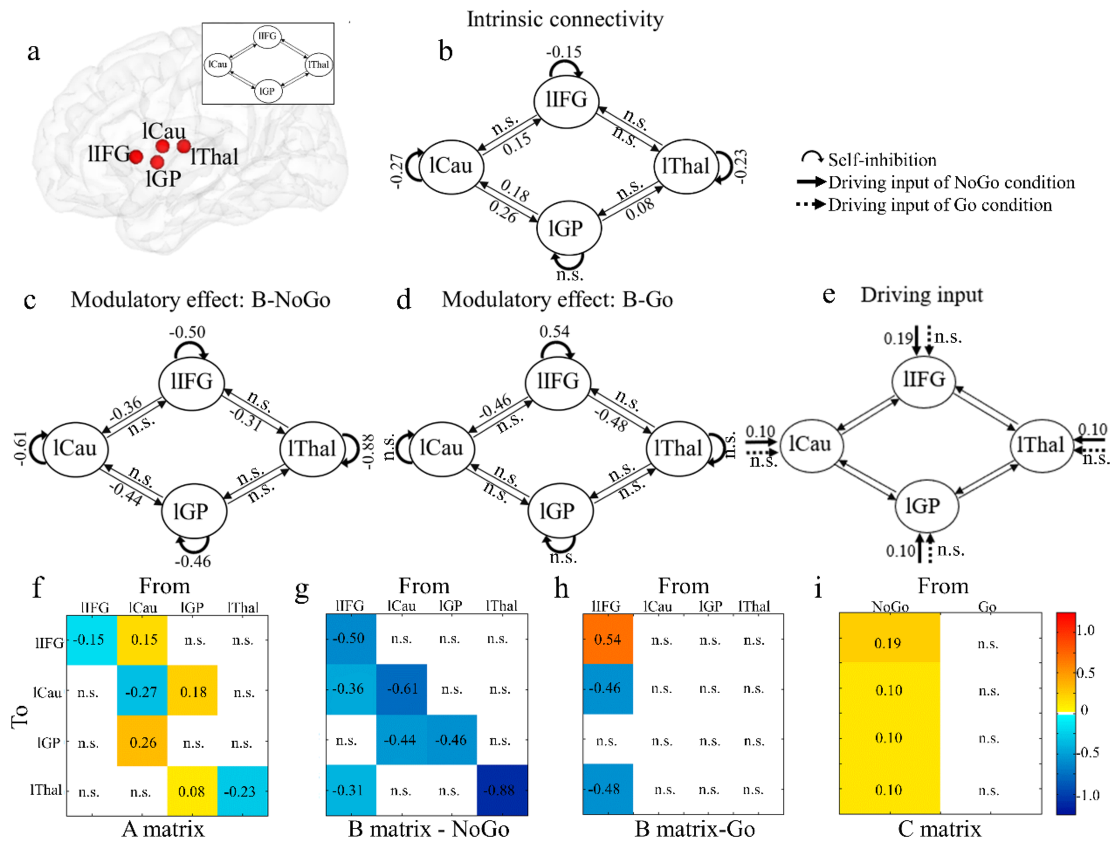


**Figure S3.** Location of regions included in the left model and group-level connectivity parameters. (a) Location of regions included in the left model. The A matrix: intrinsic connectivity independent of experimental conditions (b, f). The B matrix: modulatory effect on effective connectivity between regions and self-connections in the NoGo (c, g) and Go condition (d, h). The C matrix: driving inputs into ROIs of NoGo and Go conditions (e, i). Values in matrices reflect the connectivity parameters. Parameters with stronger evidence (posterior probability > 95%) are presented and subthreshold parameters marked with “n.s.”.
